## Supplementary Information for "Start Small: A Model for Tissue-wide Planar Cell Polarity without Morphogens"

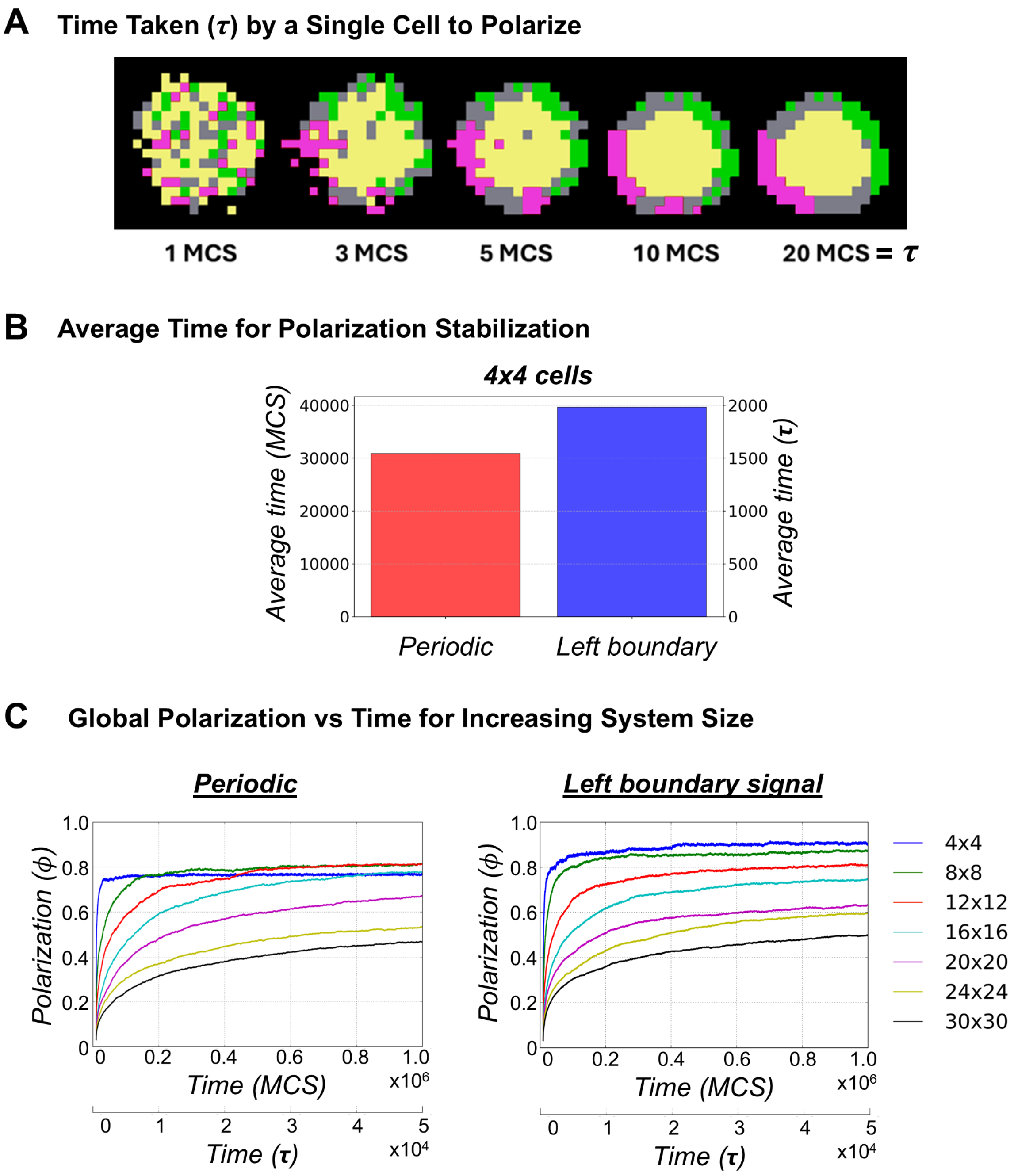

Figure S1: **Time taken for global polarization under different scenarios**
(A) An individual cell, in the absence of neighbours can establish PCP asymmetry autonomously and the domains spatially segregate in a timescale of approximately 20 MCS. This timescale is defined as the *polarization time*($\tau$) (B) Average time for polarization stabilization in 4×4 cell system for periodic boundary and left boundary signal configurations. The average time required for polarization stabilization is determined by binning the time series data and identifying the first interval where the variance in global polarization is minimal. (C) Global polarization vs. time for increasing system size for periodic boundary conditions and the configuration with left boundary signal respectively.

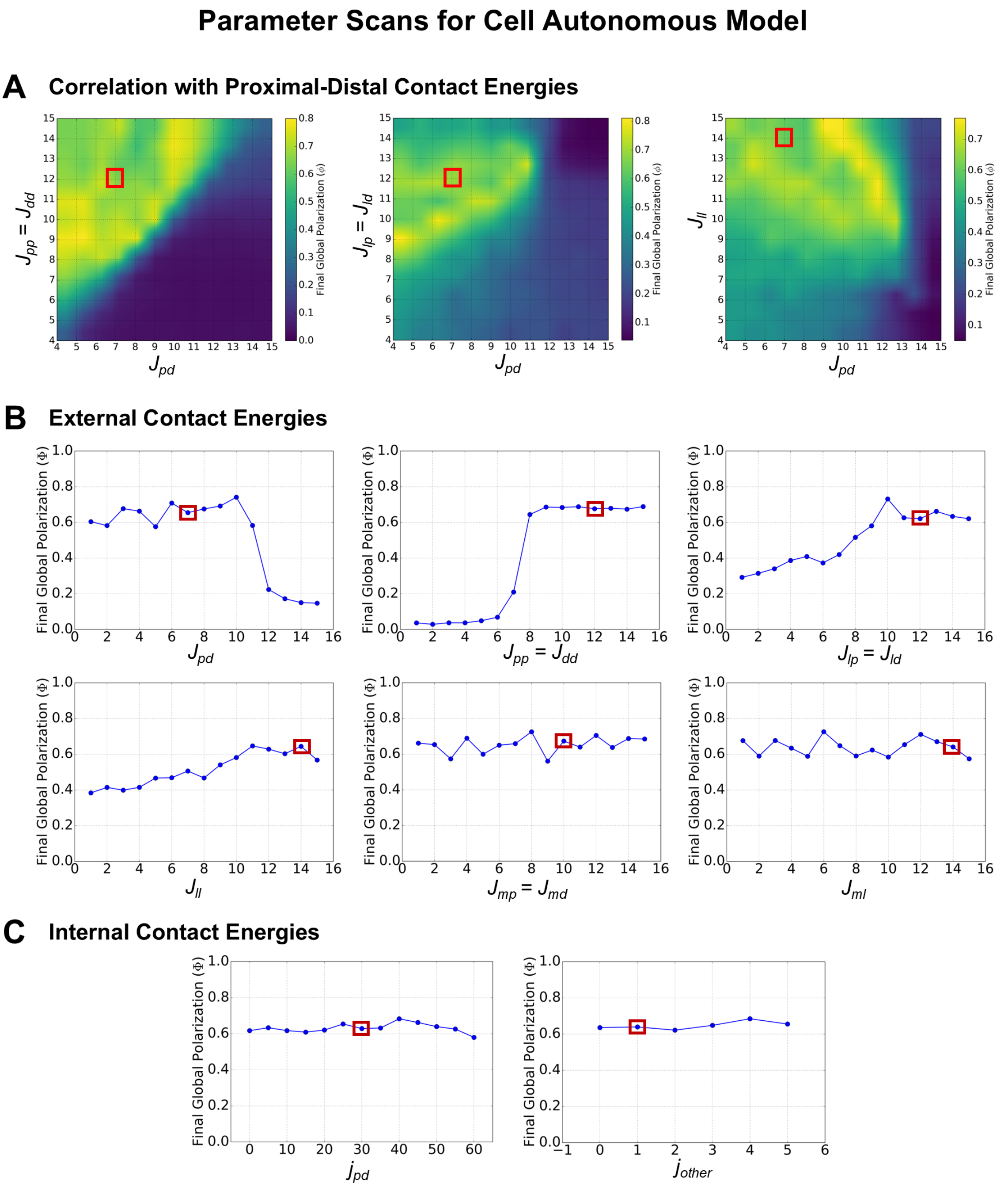

Figure S2: **Parameter sensitivity analysis of contact energies for cell autonomous model.** The final global polarization was evaluated for a $4 \times 4$ cell system under periodic boundary conditions after $10^5$ MCS. When varying one contact energy, all other energies and parameters were held fixed as listed in Tables 1, 2, and S1. (A) Correlations between the proximal-distal external contact energy ($J_{pd}$) and other external contact energies: proximal-proximal = distal-distal ($J_{pp} = J_{dd}$, left); proximal-lateral = distal-lateral ($J_{lp} = J_{ld}$, middle); and lateral-lateral ($J_{ll}$, right) (B) Sensitivity of global polarization to variations in each external contact energy, tested one at a time in the following order: $J_{pd}$, $J_{pp} = J_{dd}$, $J_{lp} = J_{ld}$, $J_{ll}$, $J_{mp} = J_{md}$, and $J_{ml}$. (C) Sensitivity of global polarization to variations in internal contact energies: $j_{pd}$ (proximal-distal internal contacts) and $j_{other}$ (all other internal contacts excluding proximal-distal). In all panels, the red box indicates the value used for simulations in the rest of the paper.

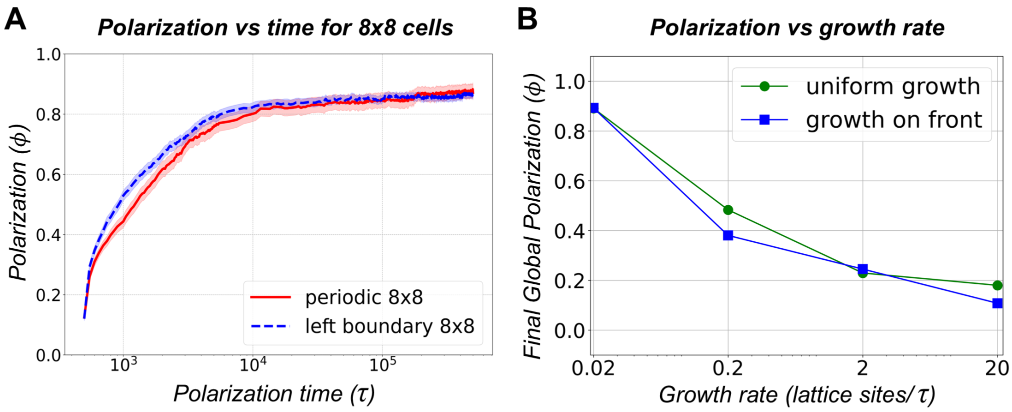

Figure S3: (A) Time evolution of global polarization, quantified by the scalar order parameter $\phi$, under periodic (red) and left-boundary (blue dashed) conditions. In both periodic and left boundary configurations, global polarization increases with increasing time in a system of $8 \times 8$ cells. The time axis is rescaled in terms of *polarization time* $\tau$. (B) Comparison of final global polarization when number of cells in the system is 900 for varying growth rates for front/distal proliferation (blue) and uniform cell proliferation (green). For uniform cell proliferation, the refractory time between cell divisions is set to $10^4$ MCS ($500\tau$). Growth rate is indicated in the units of lattice sites/$\tau$.

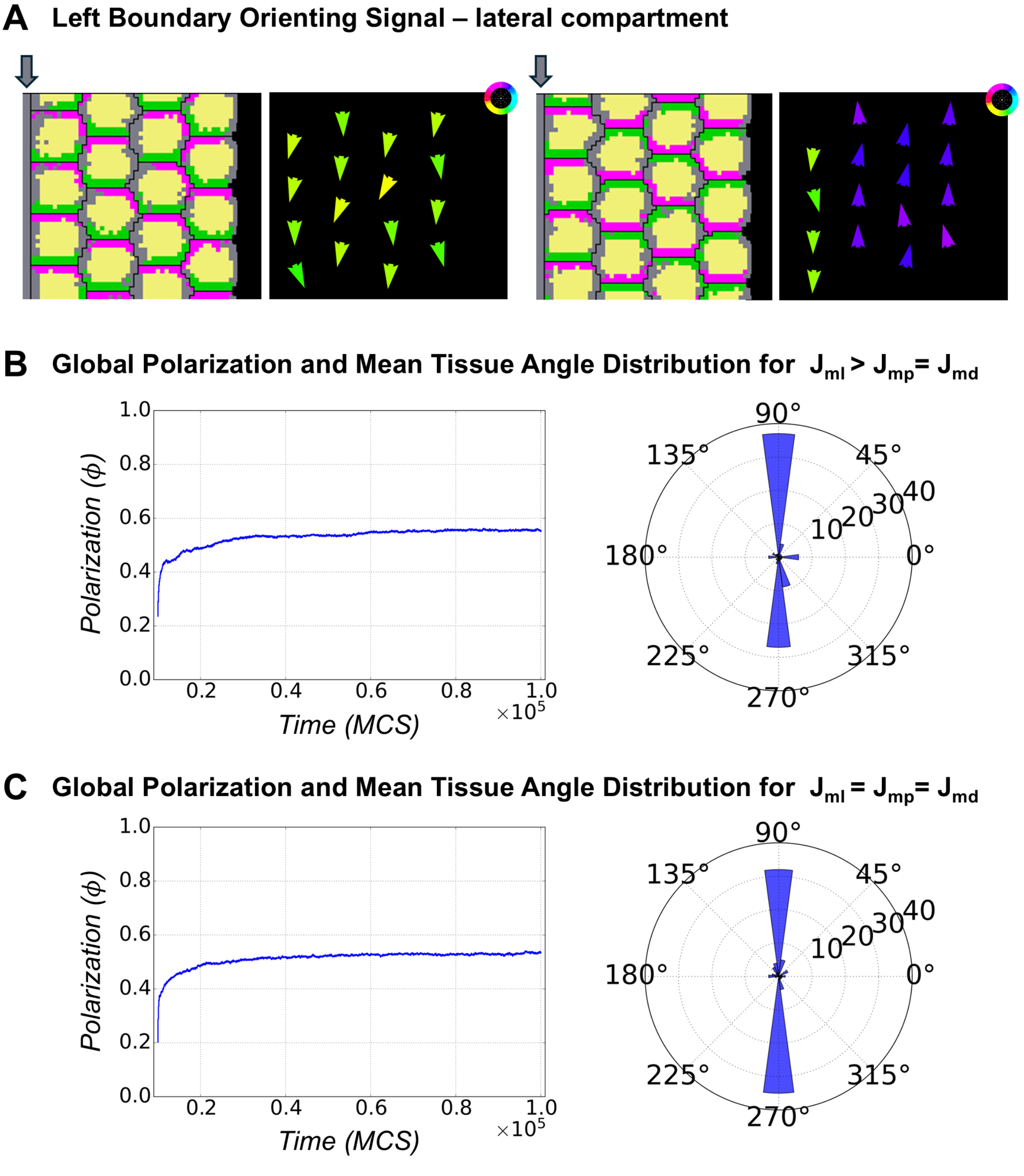

Figure S4: **Left boundary orienting signal of lateral cell type.**
(A) The left boundary orienting signal with the adhesion properties of lateral cell compartment is added. Two of the possible outcomes are shown- all cells point down (left panel), first column of cells points down whereas other columns point up. Global polarization (left panel) and mean tissue angle distribution (right panel) for (B) Higher preference of the Medium for proximal or distal compartments over the lateral compartment ($J_{ml} > J_{mp} = J_{md}$). (C) Equal preference of the Medium for proximal, distal, and lateral compartments ($J_{ml} = J_{mp} = J_{md}$). Cells do not align along the proximal-distal axis; instead, each column of cells independently chooses between up/down alignment. As a result, the average global polarization decreases, and the mean tissue polarization angle is biased around $\pm 90°$.

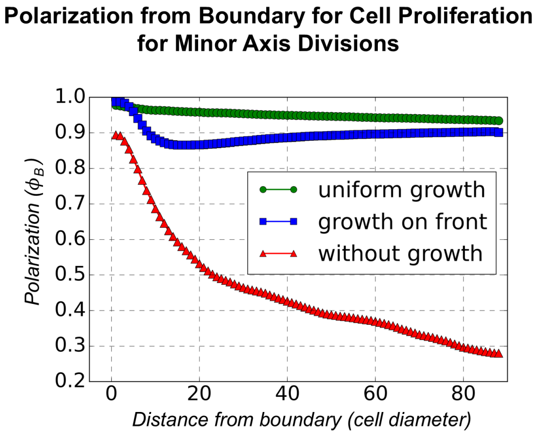

Figure S5: **Polarization from boundary for cell proliferation for minor axis divisions**
Local polarization decreases with increasing distance from the boundary but stabilizes after approximately 20 columns of cells, for systems with cell proliferation (both uniform cell proliferation and proliferation on front). However, for the system without cell proliferation, the local polarization decreases with increasing distance from the boundary.

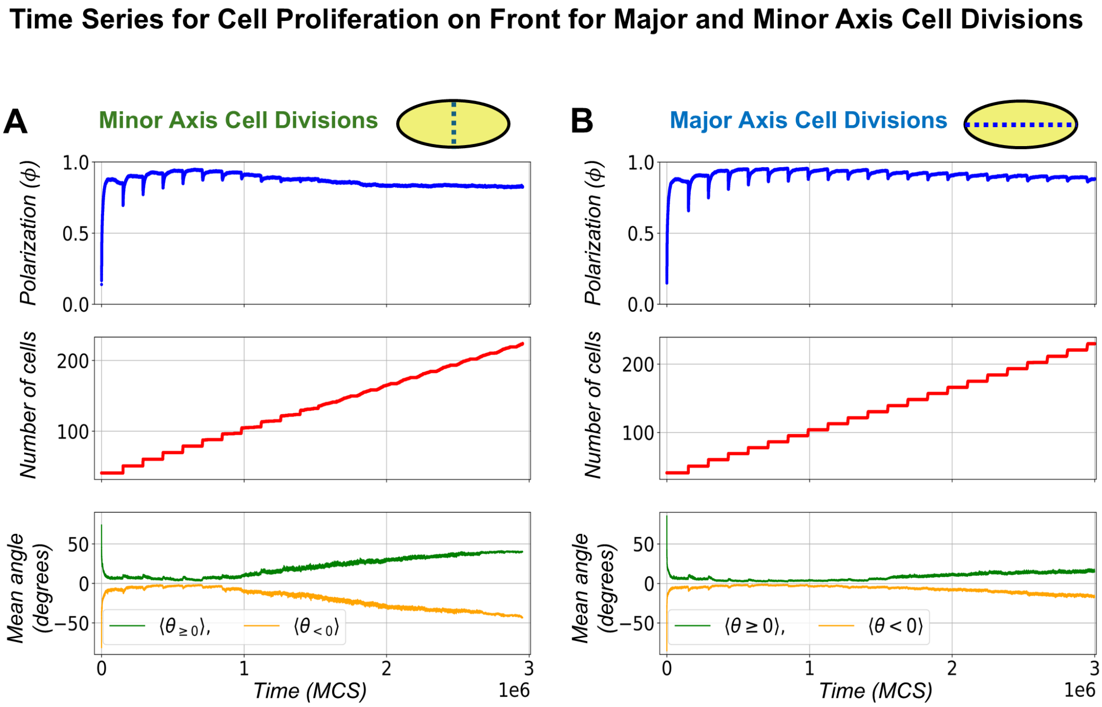

Figure S6: **Time series data of global polarization, total number of cells and the mean angle of tissue polarization for cell proliferation at the front boundary for (A) minor and (B) major axis cell divisions.** The mean polarization angle is calculated separately for positive and negative values. The mean angle deviates from 0° to ±50° as the global polarization starts decreasing.

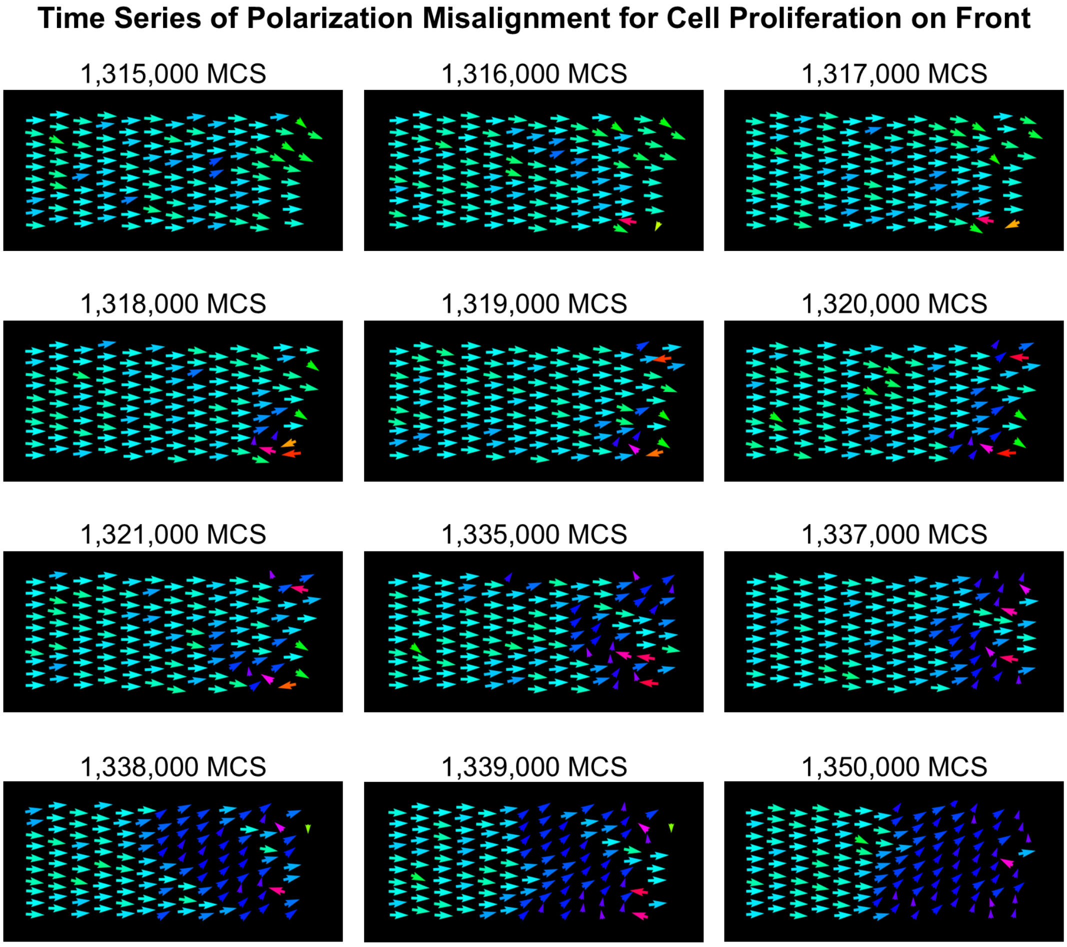

Figure S7: **Time Series of Polarization Misalignment for Cell Proliferation on Front**
Initially, all cells are aligned along the proximal-distal axis. Over time, a defect emerges in one or more cells at the front boundary, which back-propagates through the tissue, eventually leading to a change in the overall alignment direction for the cells at the front boundary. However, cells near the left boundary signal retain their alignment along the proximal-distal axis.

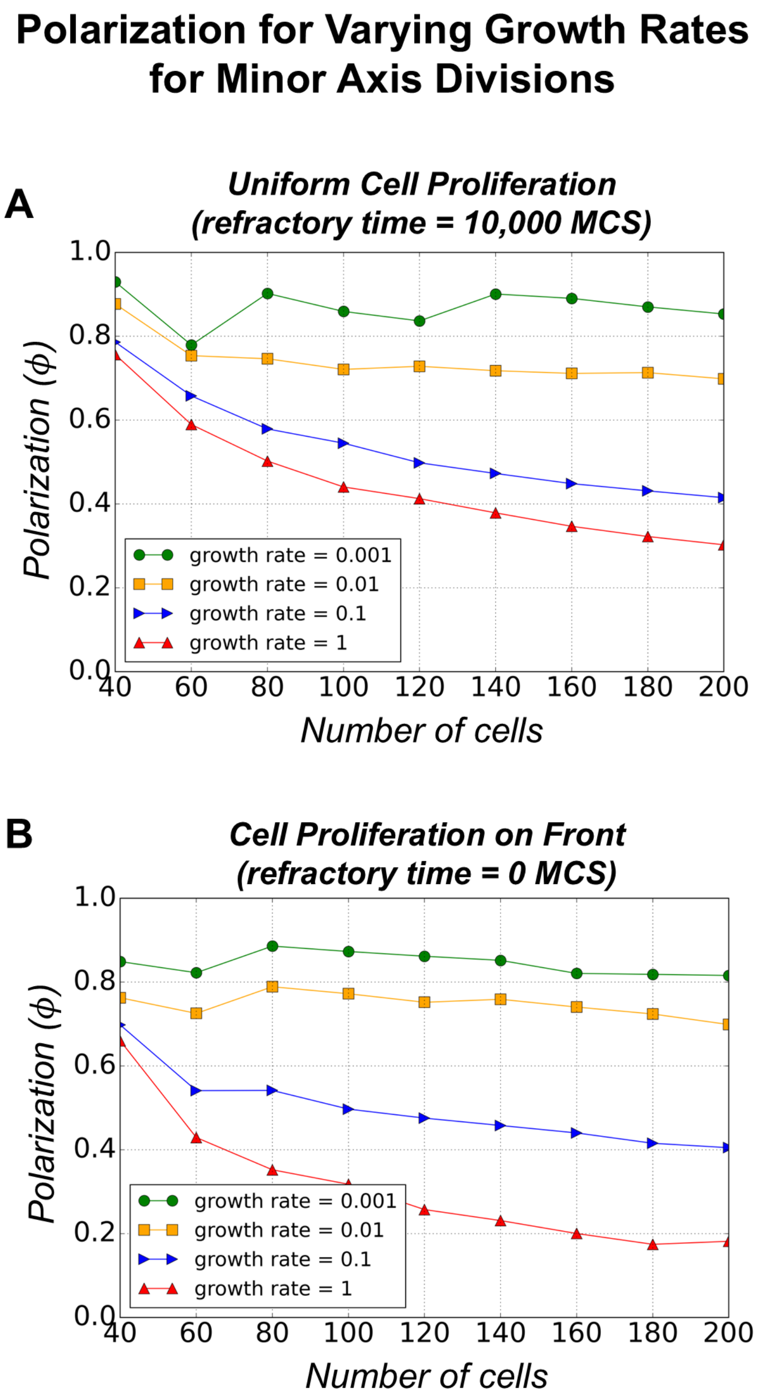

Figure S8: **Comparison of polarization for varying growth rates for cell proliferation (minor axis cell divisions).**
Comparison of global polarization vs number of cells for different growth rates under two proliferation scenarios. (A) Uniform cell proliferation with cell division along minor axis and refractory time=$10,000$ MCS. Global polarization decreases with increase in growth rate of cells for all system sizes. There is a decrease in global polarization with increase in the number of cells in the system independent of growth rate. (B) Cell proliferation on front with cell division along minor axis. Global polarization decreases with increase in growth rate of cells for all system sizes. Also, there is a decrease in global polarization as the system size increases independent of growth rate.

### Effects of Refractory Times for Uniform Cell Proliferation

#### A Tissue polarization

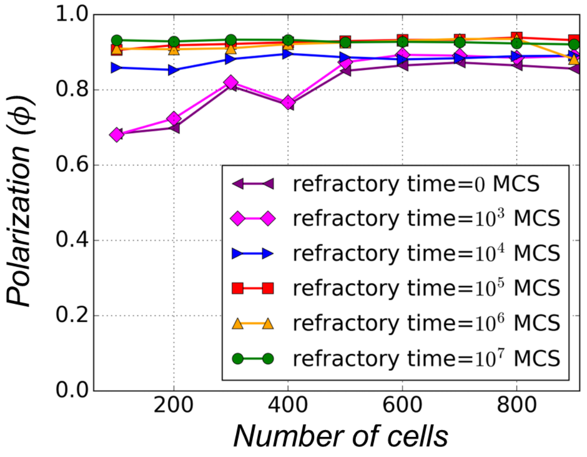

#### B Mean Angle of Polarization

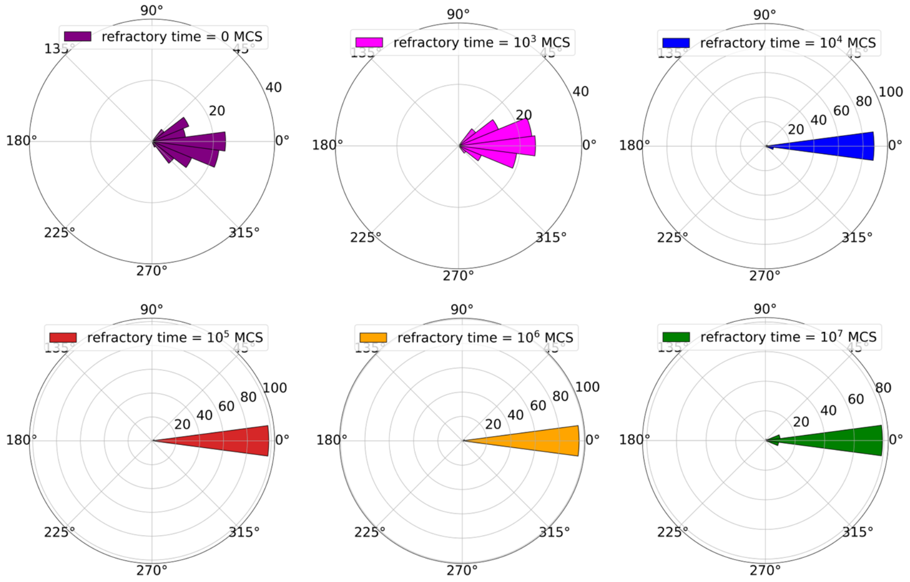

Figure S9: **Comparison of polarization and the mean angle of polarization for uniform cell proliferation (along minor axis) for varying refractory times.**
The refractory time ($t_{\text{refractory}}$) is defined as the maximum interval after a cell divides during which it will begin growing again. Specifically, after a cell divides at time $t_{\text{divide}}$, the next growth event is initiated at a time randomly chosen from the interval $[t_{\text{divide}}, t_{\text{refractory}}]$. (A) Final global polarization and (B) mean angle of tissue polarization are compared for varying refractory times ($t_{refractory}$) with increasing number of cells in the system. The global polarization remains largely unchanged as the refractory time decreases, except at $t_{\text{refractory}} = 10^4$ MCS, $10^3$ MCS, and 0 MCS where a slight reduction is observed for all system sizes. The mean polarization angle remains aligned along the proximal-distal axis in all cases except at $t_{\text{refractory}} = 10^3$ MCS, and 0 MCS where there is a small dispersion about proximal-distal axis. In all cases, cells divide along minor axis.

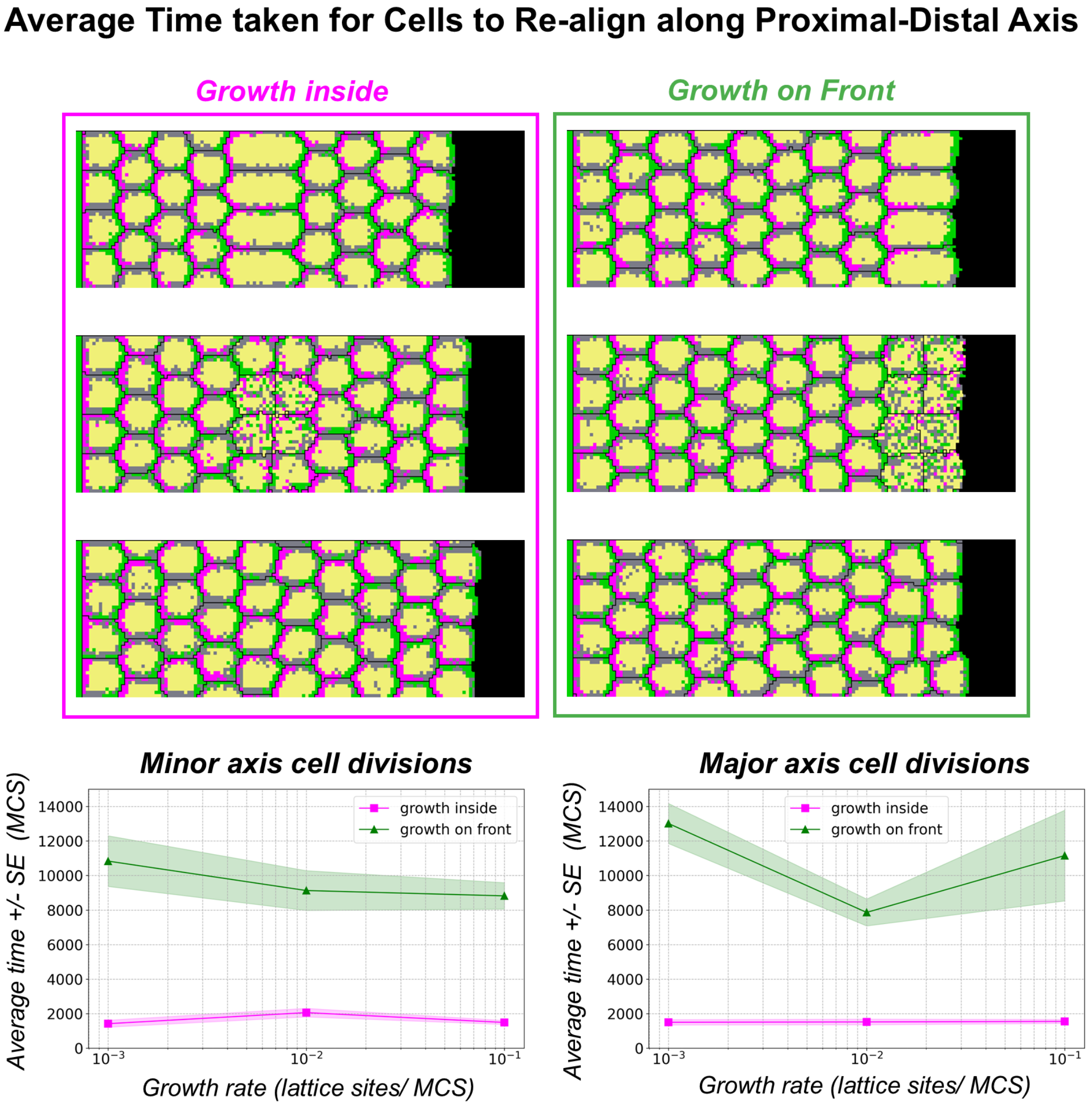

Figure S10: **Time taken by divided cells to re-polarize along proximal-distal axis for different proliferation scenarios**
Starting with a group of 4x9 cells that is already polarized, (1) the first column of cells facing the open boundary is allowed to grow and divide once (2) the fifth column of cells (fully surrounded by other cells) is allowed to grow and divide once. We then calculated the time taken for these cells to polarize along the proximal-distal axis right after they divided. This was done for an ensemble of 100 simulations and the average is plotted below for (1) growth inside and (2) growth on distal front. The results show that cells on the front take much longer to align with the rest of the tissue compared to cells that divide inside the tissue irrespective of the cleavage plane/axis of division.

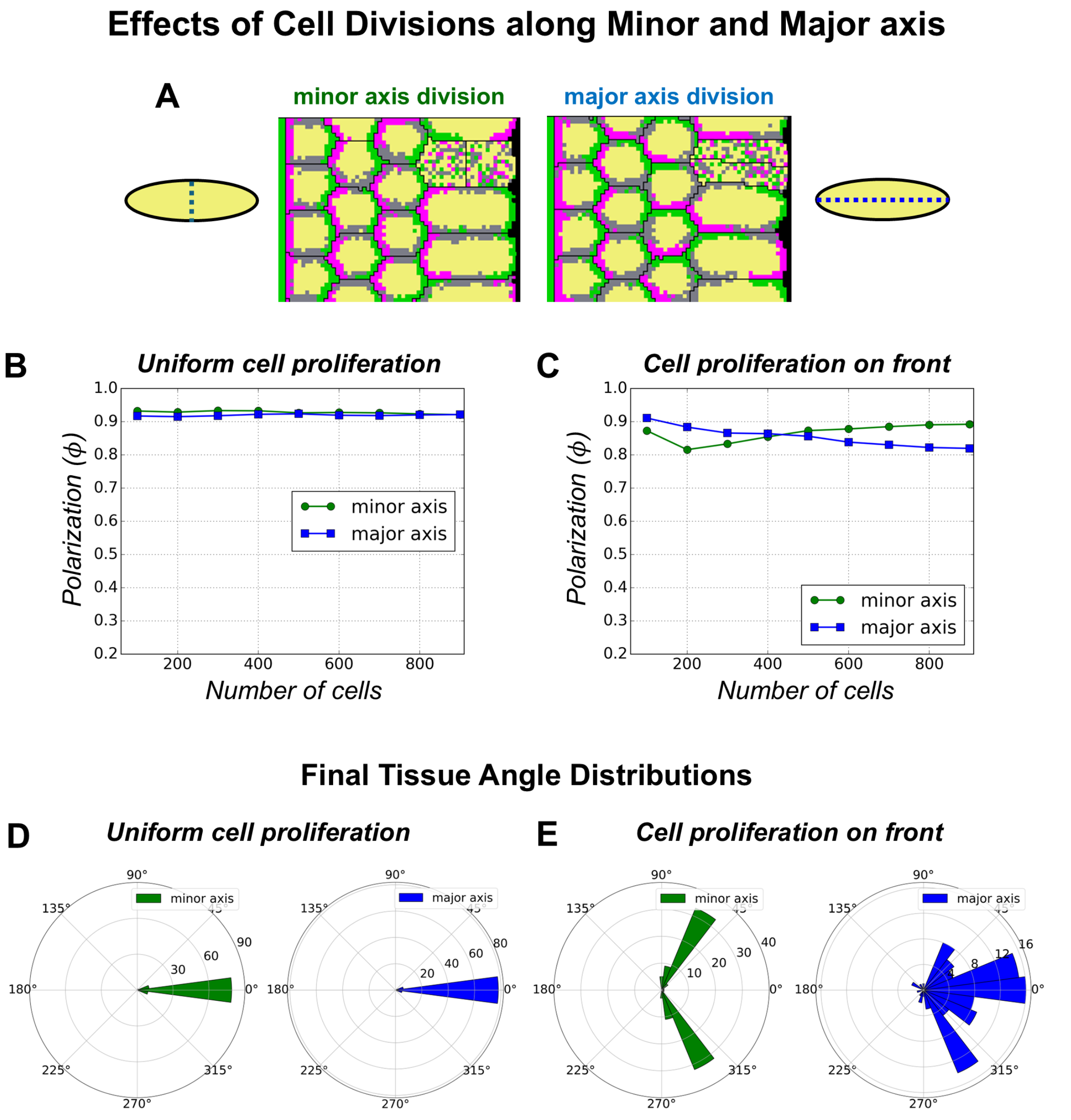

Figure S11: **Effect of the axis/plane of cell division on tissue polarization under different proliferation scenarios.**
Comparison of global polarization for cell division along minor axis (green) and major axis (blue) of the dividing cells. (A) Illustration of division planes: minor axis corresponds to the short axis of the cell, while major axis corresponds to the long axis. (B) Comparison of global polarization versus number of cells for uniform cell proliferation. The refractory time for uniform proliferation is $10^7$ MCS. The global polarization remains unchanged with change in the plane/axis of cell division. (C) Comparison of global polarization versus number of cells for proliferation on front boundary. The global polarization changes only by $\approx 1\%$ when the plane/axis of division is changed. The growth rate is 0.001 lattice sites/MCS for both proliferation scenarios. Final tissue angle distributions for (D) Uniform cell proliferation for minor (green, left panel) and major (blue, right panel) axis divisions (D) Cell proliferation on front for minor (green, left panel) and major (blue, right panel) axis divisions. The angle distribution is primarily along proximal-distal axis for uniform cell proliferation for both minor and major axis cell divisions. However, for proliferation of front scenario, the tissue angle is distributed along $\pm 60^\circ$ for minor axis divisions, whereas the angle is dispersed about the proximal-distal axis for major axis divisions.

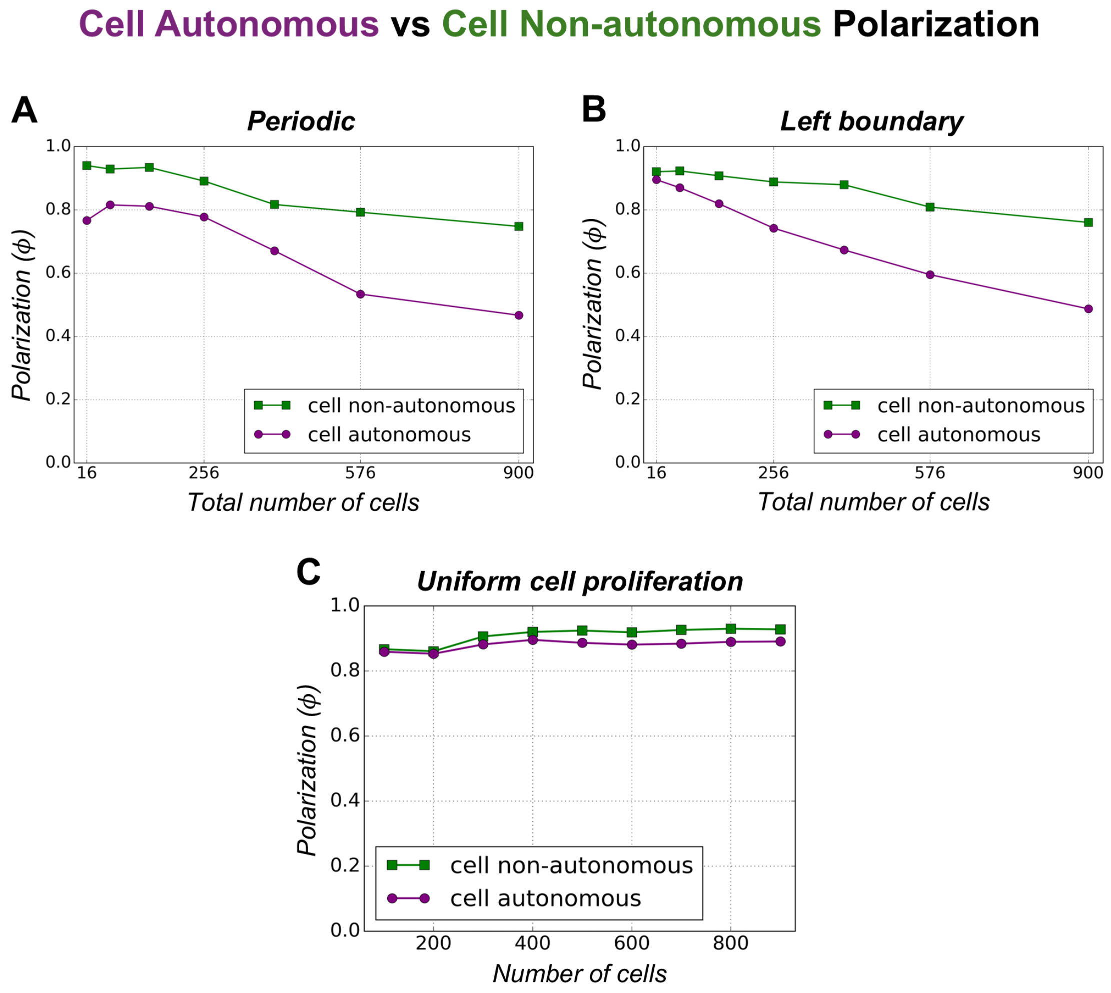

Figure S12: **Comparison of cell-autonomous vs. cell non-autonomous global polarization.**
Panel (A) shows the comparison of global polarization for cell-autonomous vs. cell non-autonomous global polarization under periodic boundary conditions with increasing numbers of cells (final state after $10^6$ MCS). Panel (B) shows the same comparison under left boundary signal configurations. Panel (C) shows the comparison of global polarization for cell-autonomous vs. cell non-autonomous with increasing number of cells for uniform cell proliferation with a refractory time of 10,000 MCS and a growth rate of 0.001 lattice sites/MCS.

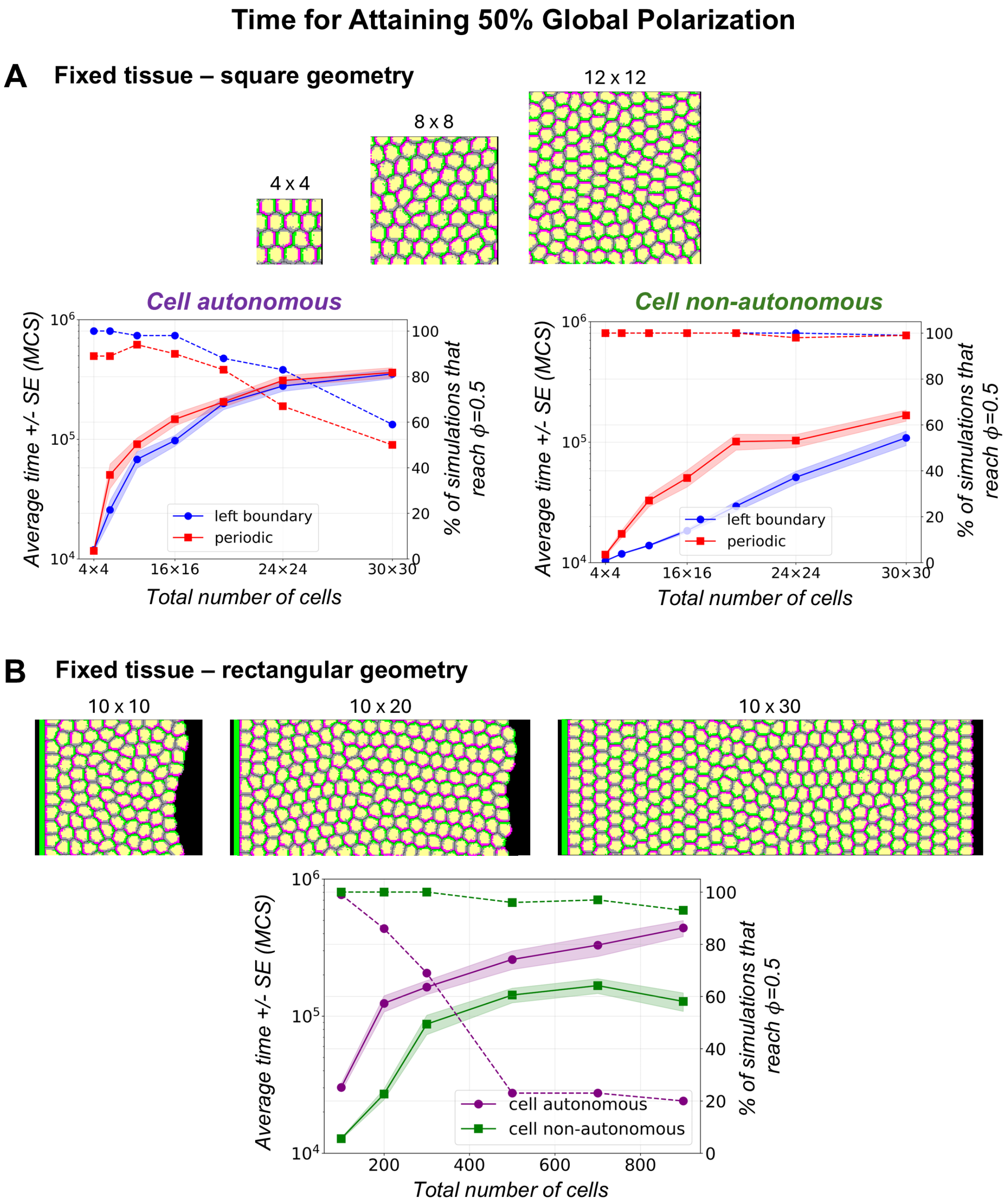

Figure S13: **Comparison of average time taken to reach 50% global polarization for cell-autonomous and cell non-autonomous polarization scenarios.**
(A) **Cell-autonomous polarization** (left): As system size increases along both $x$ and $y$ directions (square geometry) under periodic boundary and left boundary signal configurations, the time to attain 50% global polarization (solid line) increases, and the fraction of simulations that reach 50% global polarization (dashed line) decreases by approximately 50%. (right) **Cell non-autonomous polarization**: As system size increases under the same conditions, the time to reach 50% global polarization (solid line) also increases, but less than in the cell-autonomous case, while the fraction of simulations reaching 50% polarization (dashed line) remains roughly constant near 100%. (B) With system size increased along $x$ direction while keeping 10 cells constant along $y$ direction (rectangular geometry), the time to reach 50% global polarization (solid line) increases with increasing system size for both autonomous and non-autonomous cases, but the increase is higher for cell-autonomous. Although the fraction of simulations reaching 50% polarization (dashed line) decreases with system size for both cases, the decrease is very small for non-autonomous cells, whereas it drops sharply to 20% for cell-autonomous cells.

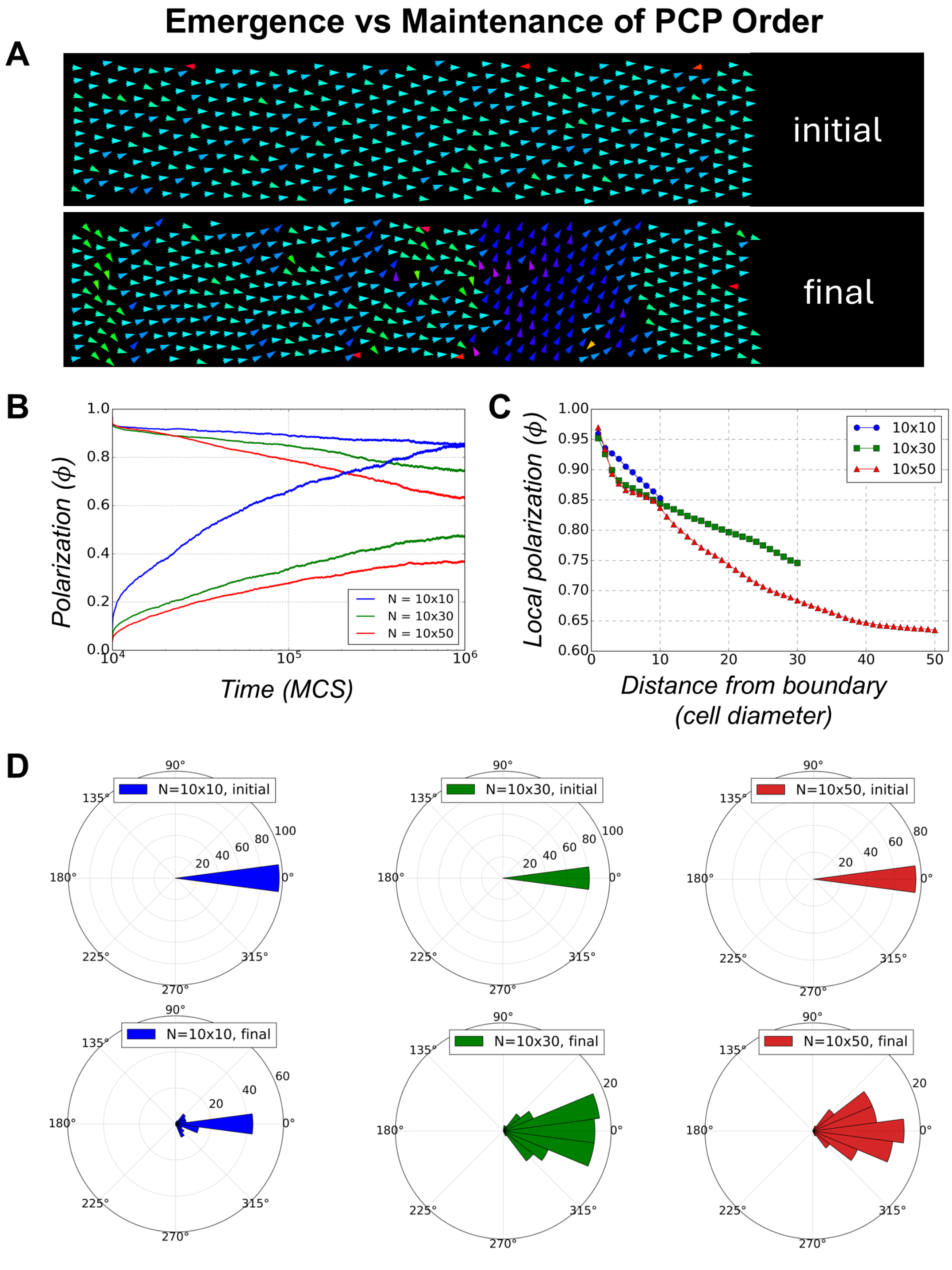

Figure S14: **Emergence vs maintenance of PCP order**
The maintenance of a pre-established PCP order is studied for a system where **seed** cells fill up the lattice and realx their boundaries. Then, the proximal and distal compartments are allocated to left and right ends of each of seed cells. The maintenance of this polarity is evaluated as time progress until $10^6$ MCS. (A) The initial and final polarity vectors for a representative simulation are shown. (B) Comparison of global polarization and local polarization for increasing system sizes for systems with pre-established PCP order and systems where PCP is emerging. Global polarization decreases linearly for systems that start with PCP order whereas it increases for systems where PCP order emerges from a random initial configuration. (C) As the distance from the left boundary increases, the local order decreases for increasing system size. (D) Initial (top panels) and final (bottom panels) tissue angle distributions for increasing system sizes ($10 \times 10$- left panel, blue; $10 \times 30$- middle panel, green; $10 \times 50$- left panel, red). As system size increases, the tissue angle is more dispersed about the proximal-distal axis with increasing time.

#### Emergence vs Maintenance of PCP Order for Perfect Hexagonal Packing

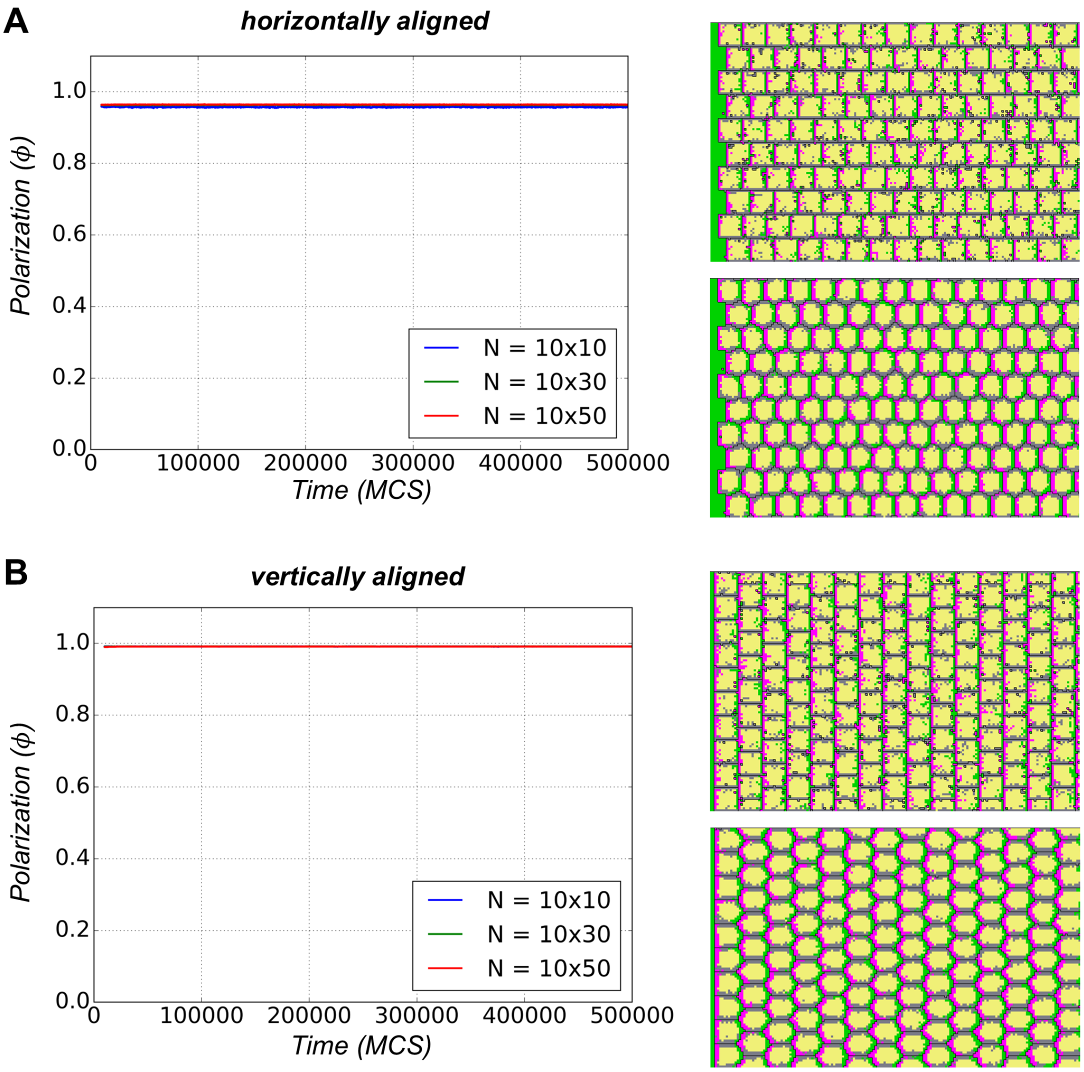

Figure S15: **Emergence vs maintenance of PCP order for perfect hexagonal packing**
The maintenance of a pre-established PCP order is studied for a system with perfect hexagonal packing for (A) horizontal and (B) vertical alignment.

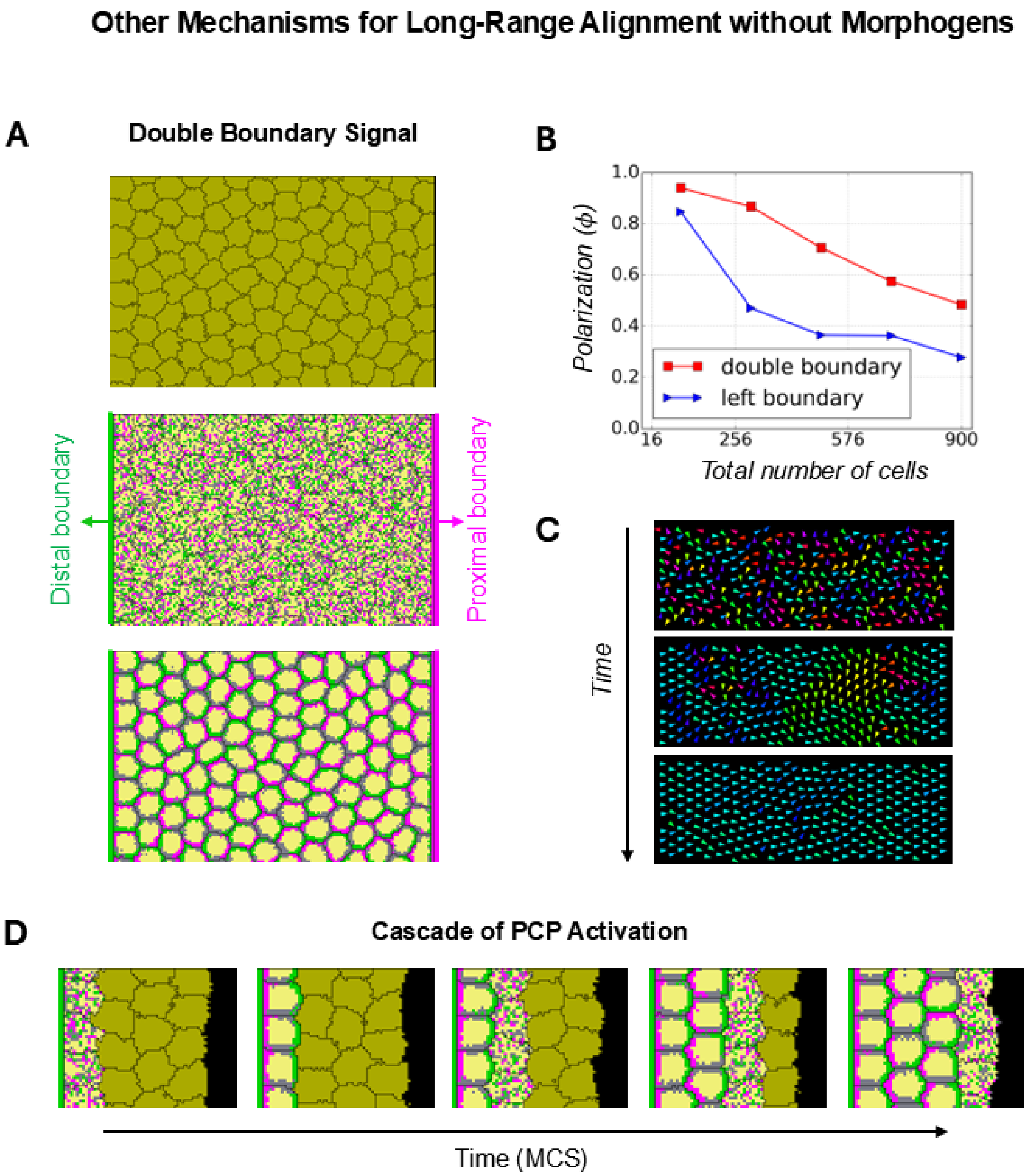

Figure S16: **Other mechanisms for long range alignment without morphogens.** **(A) Double Boundary Signal.** Schematic of the double boundary signal configuration with complementary signalling boundaries on opposite sides of the tissue. (B) Global polarization for the double boundary configuration across system sizes, showing increased polarization (red) compared to the single left boundary case (blue). (C) Simulation snapshots of a $10 \times 30$ cell system polarized along the proximal–distal axis by double boundary signals over time. **(D) Cascade of PCP Activation.** Cell columns are sequentially activated for PCP signalling from a local boundary.

Table S1: **Simulation Parameters:** This table lists all parameters used in the simulations, including volume constraints, neighbour order for lattice site copy events and contact energy calculations and volume fractions of each cell and cell compartments inside the cell.

| **Parameter** | **Name** | **Value** |
|---|---|---|
| $T$ | CPM fluctuation amplitude | 15 |
| $n_{\text{copy}}$ | Neighbor range for lattice site copy attempts | 3 |
| $n_{\text{contact}}$ | Neighbor range for contact energy calculations | 4 |
| $t$ | Total time (depends on the case examined) | $10^5$ to $5 \times 10^7$ MCS |
| $cd$ | Cell Diameter | 12 |
| $tV$ | Target Volume | 144 |
| $\lambda_{\text{v}}$ | Strength of Volume Constraint | 12 |
| **Volume Fractions** | | |
| $p_{\text{P}}$ | Proximal Domain | 0.15 |
| $p_{\text{D}}$ | Distal Domain | 0.15 |
| $p_{\text{C}}$ | Cytoplasmic Domain | 0.5 |
| $p_{\text{L}}$ | Lateral Domain | 0.2 |
| **Cell Proliferation Parameters** | | |
| growth_rate | growth rate for proliferation | 0.001 |
| $t_{\text{relax}}$ | relaxation time for cell proliferation | varies from 0 to $10^7$ MCS for uniform cell proliferation |

#### Supplementary Movies

Movie 1: **Time Evolution of a system of 8×8 cells under periodic boundary conditions**
Time evolution of an 8×8 cell system with periodic boundary conditions over $10^5$ Monte Carlo steps (MCS). Vector fields show the evolution of the cell polarity vectors throughout the simulation. The final global polarization for this simulation is 0.75.

Movie 2: **Time Evolution of a system of 30×30 cells under periodic boundary conditions**
Time evolution of a 30×30 cell system with periodic boundary conditions over $10^5$ Monte Carlo steps (MCS). Vector fields show the evolution of the cell polarity vectors throughout the simulation. Domains of local alignment and swirling patterns can be seen. The final global polarization for this simulation is 0.28.

Movie 3: **Time Evolution of a system of 8×8 cells under left boundary signal configuration**
Time evolution of an 8×8 cell system with left boundary signal over $10^5$ Monte Carlo steps (MCS). Vector fields show the evolution of the cell polarity vectors throughout the simulation. All cells align along proximal-distal axis. The final global polarization for this simulation is 0.94.

Movie 4: **Time Evolution of a system of 30×30 cells under left boundary signal configuration**
Time evolution of a 30×30 cell system with left boundary signal over $10^5$ Monte Carlo steps (MCS). Vector fields show the evolution of the cell polarity vectors throughout the simulation. Cells align along proximal-distal axis closer to the boundary and alignment is lost after a few columns from the left boundary. The final global polarization for this simulation is 0.35.

Movie 5: **Time Evolution of a system under proliferation on front boundary**
Starting from a configuration of 10×4 cells polarized along the proximal–distal axis by a left boundary signal, cells at the front boundary grow and divide. After several rounds of division, alignment along the proximal–distal axis is lost. The simulation runs for a total of $10^6$ Monte Carlo steps (MCS), during which the number of cells increases to 105 and the global polarization reaches 0.84. Vector fields depict the evolution of cell polarity vectors throughout the simulation.

Movie 6: **Time Evolution of a system under uniform proliferation**
Starting from a configuration with 10x4 cells (which polarize along proximal-distal axis with the help of a left boundary signal), all cells in the tissue grow and divide (cells start to grow at random times in the interval $[0, 10^5]$ MCS. After a few rounds of division, the alignment along proximal-distal axis is lost. The simulation runs for a total of $10^6$ Monte Carlo steps (MCS), during which the number of cells increases to 522 and the global polarization reaches 0.95. Vector fields show the evolution of the cell polarity vectors throughout the simulation.
